# Bernstein polynomial approximation of fixation probability in finite population evolutionary games

**DOI:** 10.1101/2022.08.05.502960

**Authors:** Jiyeon Park, Paul K. Newton

**Author notes:** Email addresses:* (Jiyeon Park), (Paul K. Newton).

## Abstract

We use the Bernstein polynomials of degree *d* as the basis for constructing a uniform approximation to the rate of evolution (related to the fixation probability) of a species in a two-component finite-population frequency-dependent evolutionary game setting. The approximation is valid over the full range 0 ≤ *w* ≤ 1, where *w* is the selection pressure parameter, and converges uniformly to the exact solution as *d* → ∞. We compare it to a widely used non-uniform approximation formula in the weak-selection limit (*w* ∼ 0) as well as numerically computed values of the exact solution. Because of a boundary layer that occurs in the weak-selection limit, the Bernstein polynomial method is more efficient at approximating the rate of evolution in the strong selection region (*w* ∼ 1) (requiring the use of fewer modes to obtain the same level of accuracy) than in the weak selection regime.

## 1. Introduction

In finite-population stochastic evolutionary games, an important quantity is the so-called fixation probability of a given sub-species of *mutants* [1]. For a population of size *N*, we denote this fixation probability for a sub-species *A* comprised of *i < N* mutants as 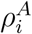, and note that it is related to the rate of evolution of that sub-species via 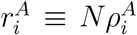. It is straighforward to show that the fixation probability of a sub-species, in the absence of selection, is simply given by the neutral drift formula 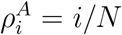 [2]. With rate of evolution 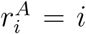 this says that any mutant has an equal probability of reaching fixation (spreading throughout the population), and the rate of evolution is simply given by the number of mutants. When selection is present, sometimes introduced using a selection pressure parameter 0 ≤ *w* ≤ 1, the fixation probability formulas 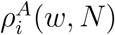 are continuous functions of this parameter, as well as *N*, and become more complicated. Of particular interest are the *weak selection* (individuals have very similar fitness) and *strong selection* (individuals have widely varying fitness) limits, given respectively by *w <<* 1 and *w* ∼ 1. In those two cases, asymptotic approximations or Taylor expansions are often invoked [3]. However, since these formulas are simultaneously functions of *N*, they are local approximations (in *w*) only, and are interpreted as being valid for fixed population sizes *N*. As simultaneous functions of *w* and *N*, however, these approximations break down, particularly in the small (*wN <<* 1) and large (*wN >>* 1) *population* limits, or intermediate values of the selection pressure parameter *w*. That is because in these two-parameter formulas, these limits become singular, and contain boundary layers (non-uniformities), rendering them invalid throughout the full selection range [4] and for all population sizes *N*.

In this note, we take a different approach and derive a new family of formulas, based on the Bernstein polynomial basis set [5]. These can be used as global approximation formulas in the full selection interval 0 ≤ *w* ≤ 1, for all *N*, that uniformly approximate the more complicated exact fixation probability and rate of evolution formulas. Bernstein polynomials were originally constructed by S. Bernstein [5] in 1912 and used to prove the Weierstrass approximation theorem [6] which states that any continuous function on a closed interval can be uniformly approximated by a series of polynomials. They have since become a standard tool in the field of polynomial approximation theory [6, 7, 8]. If *B*_*d*_(*r*_*i*_; *w*) denotes the Bernstein polynomial (of degree *d*) representation of the rate of evolution *r*_*i*_, we will show that for each *i, B*_*d*_(*r*_*i*_; *w*) uniformly converges [6] to *r*_*i*_ as *d* → ∞ in the full interval *w* ∈ [0, 1]. We investigate properties of the representation for different selection regimes *w* ∼ 0 and *w* ∼ 1 and values of *d* since we anticipate that low *d* representations are more useful in practice.

## 2. The model

### 2.1. The finite-population evolutionary game

We consider a frequency-dependent Moran process model [1, 3, 9] comprised of *N* individuals divided into two sub-populations *A* (mutants) and *B* (wild-type). Let *i* be the number of individuals in sub-population *A*, let *j* be the number of individuals in sub-population *B* with *i* + *j* = *N*. Let 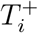 be the transition probability from state *i* to *i* + 1, and let 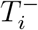 denote the transition probability from state *i* to *i* − 1. Then 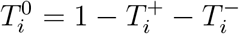 denotes the probability that the system stays at its current state (under the assumption of no mutations). Both *i* = 0 and *i* = *N* are absorbing states repectively corresponding to a homogeneous *B* population and a homogeneous *A* population, and these lead to 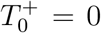 and 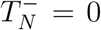, or 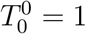 and 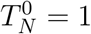. For all other states, the Markov transition probability 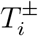 is computed by considering a weighted birth-death process [10]. The birth rates of this process are assumed to be proportional to the expected fitness of populations *A* and *B*, denoted 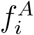 and 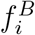, which are, in turn, functions of the expected payoffs, 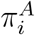 and 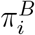:

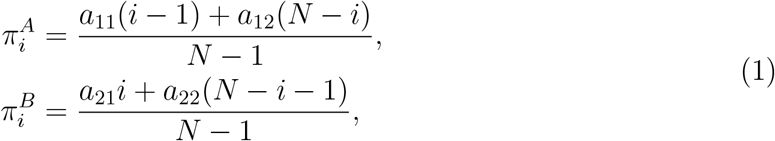

with 2 × 2 payoff matrix:

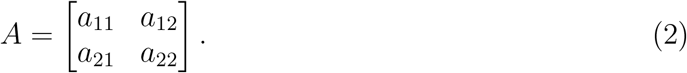

The fitness functions are defined by:

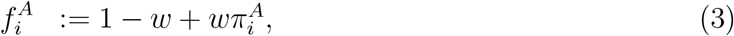

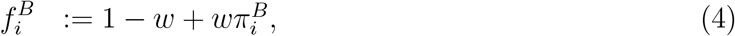

where 0 ≤ *w* ≤ 1 is selection intensity parameter, with *w <<* 1 and *w* ∼ 1 being the weak selection and strong selection limits, respectively [1, 3, 9, 11]. To compute the transition probabilities 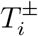, we use fitness-weighted frequency-dependent averages 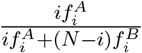 and 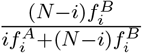 as birth rates of the *A* and *B* sub-populations respectively, which give rise to transition probabilities:

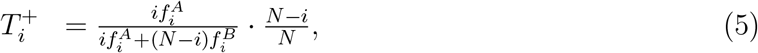

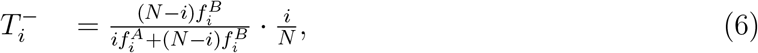

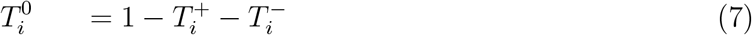

for *i* ∈ {1, …, *N* − 1}. Knowing *T*_0_ = 1 and *T*_*N*_ = 1 are two absorbing states, we can now write the (*N* + 1) × (*N* + 1) tridiagonal transition matrix *T* governing the Markov process [10]:

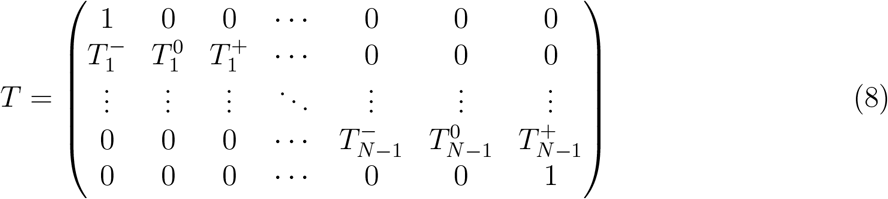

for this finite-population evolutionary game.

### 2.2. Fixation probability formulas

An important quantity for these games is called the fixation probability that a finite number of mutants of type *A* invade and take over the resident population of type *B* [1]. We will denote by *ρ*_*i*_ the probability that *i* mutants of type *A* fixate the entire population for *i* = 0, 1, …, *N*. Then it is obvious by the assumption of no mutation that *ρ*_0_ = 0 and *ρ*_*N*_ = 1. To obtain an exact formula for *ρ*_*i*_, it is easiest to start from the recurrence relation (*i* ≥ 1):

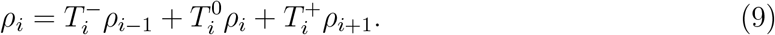

Defining variables, *ϕ*_*i*_ := *ρ*_*i*_ − *ρ*_*i*−1_ and 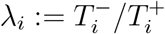, equation (9) can be re-written:

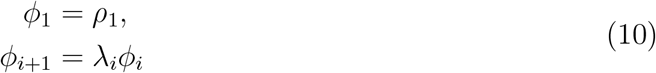

for *i* = 1, …, *N* − 1, which is equivalent to

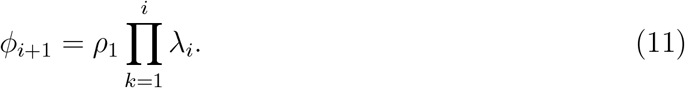

Summing (11) over *i* gives rise to 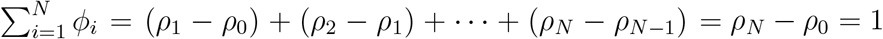 which leads to:

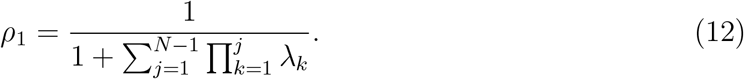

Then, using the same summation of (11) but up to any number of terms as desired, it implies 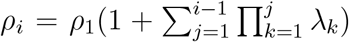, and the general fixation probability *ρ*_*i*_ of *A* starting from the state, *i*, is written as follows:

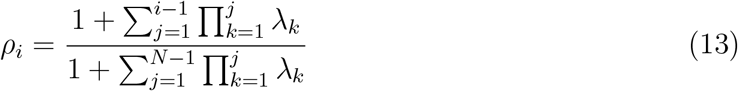

using 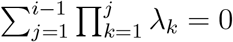 when *i* = 1. The expression in (13) is valid for any one-dimensional birth-and-death processes (without mutations) and is well known [1, 9]. Tailoring to our model, using the variable 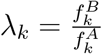, the fixation probability in (13) then becomes:

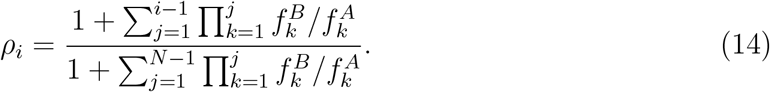

### 2.3. The repeated prisoner’s dilemma game

An approximation to the rate of evolution in the weak selection limit, 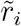, discussed in [1, 9, 11, 12], is given by:

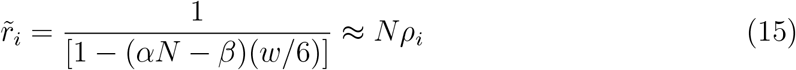

with *α* = *a*_11_ + 2*a*_12_ − *a*_21_ − 2*a*_22_ and *β* = 2*a*_11_ + *a*_12_ + *a*_21_ − 4*a*_22_ where *a*_11_, *a*_12_, *a*_21_, *a*_22_ are entries of a payoff matrix (2). It is important to keep in mind that this formula is a *local* approximation under weak selection. Due to the fact that it is a rational function of selection strength (*w*), its validity is limited to a small subinterval of [0, 1]. To appreciate the limitations of the formula as a global approximation, we consider for definiteness a repeated prisoner’s dilemma (PD) game [1, 9] in which two-players, a cooperator (*C*) and a defector (*D*), repeatedly interact with the ability to choose strategies on each interaction. The general PD payoff matrix is given by:

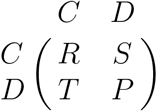

with the prisoner’s inequality

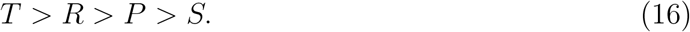

We consider a Tit-for-Tat strategy for player *A* playing against a player (*B*) who always defects. The Tit-for-Tat player cooperates in the first round (*n* = 1), then does whatever the opponent did in the previous round. If this game is repeated *n* times, it has the following payoff matrix [1, 9]

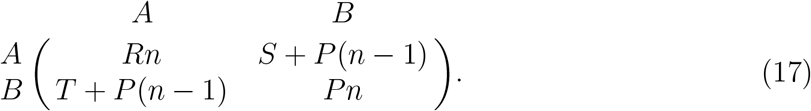

For the emergence of cooperation, we need [9] that

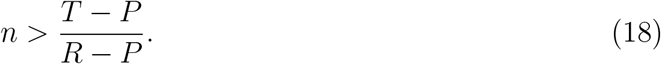

With the choice of *R* = 3, *S* = 0, *T* = 5, *P* = 1 for our experiment, (18) gives *n* ≥ 3.

The limitations of the approximation formula are shown in figure 1 where we plot the weak selection formula 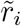 and the exact formula (14) for *N* = 10, 100, 1000, 10000 in the range *w* ∈ [0, 0.01]. Figure 1(a) shows the two formulas begin to depart with increasing *w*, while figures 1(b)-(d) show an internal boundary layer developing around the threshold value *w*_*T*_, and a negative rate of evolution when *w* is above it. The threshold value *w*_*T*_ → 0 as *N* → ∞ (i.e. the boundary layer is pushed to the left boundary). To ensure that 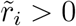, it is straightforward to obtain the following threshold formula:

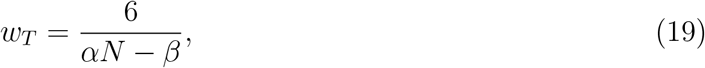

with

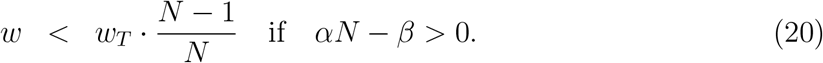

**Figure 1:**
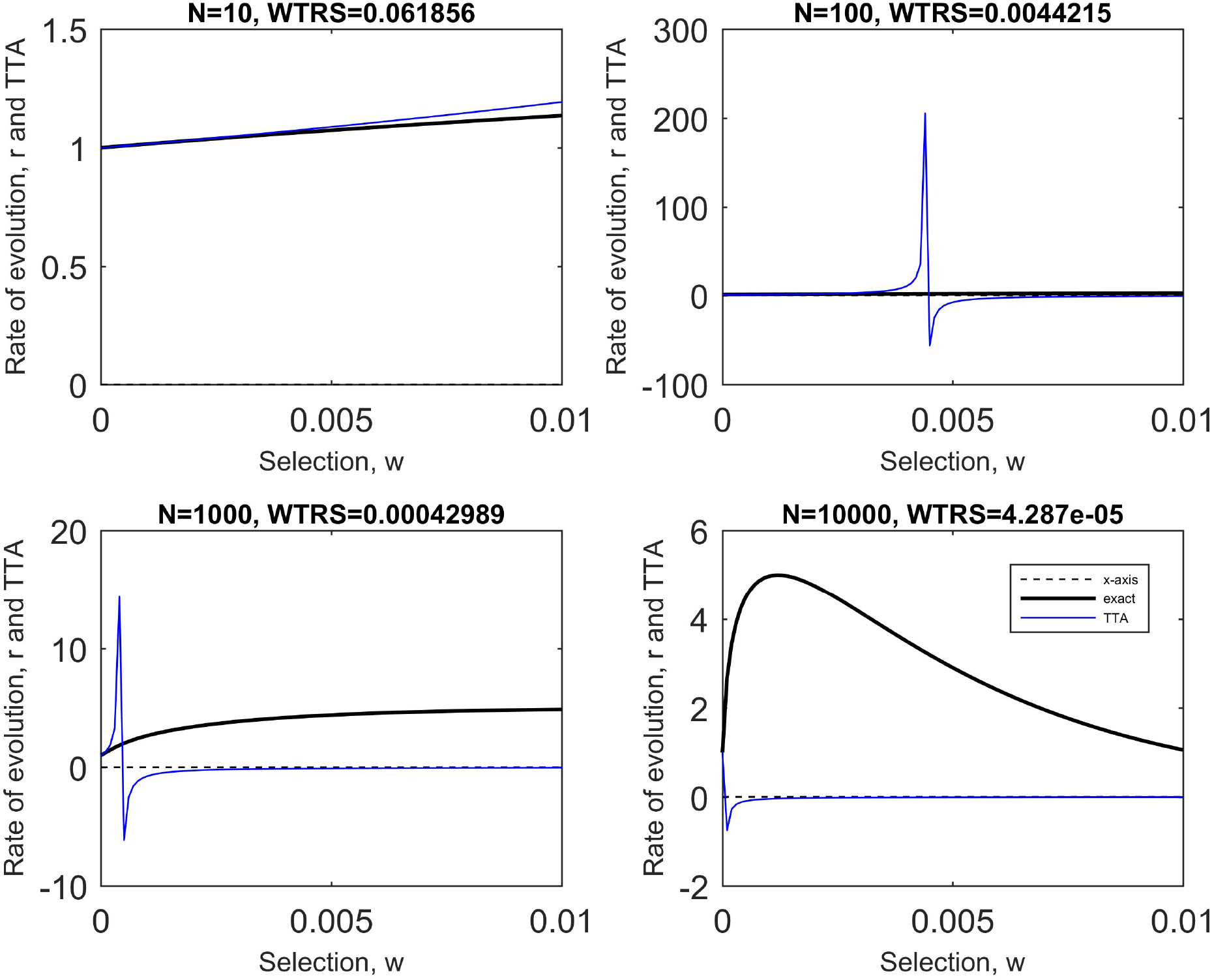
The local approximation 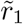 over *w* ∈ [0, 0.01]. The validity of 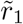 (blue line) as a local approximation to the rate of evolution *r* = *Nρ*_*A*_(black line) reduces as *N* increases since it takes nonnegative values only in the interval *w* ∈ [0, *w*_*T*_] and develops an internal boundary layer around *w*_*T*_ rendering the approximation non-uniform. We use the repeated Prisoner’s Dilemma game values *a*_21_ = 5 *> a*_11_ = 3 *> a*_22_ = 1 *> a*_12_ = 0, and *n* = 10, but the non-uniformity in the approximation is a general feature associated with the behavior of the fixation probability curve as a function of both *w* and *N*.

The convergence *w*_*T*_ → 0 as *N* → ∞ is shown in figure 1.

## 3. Bernstein polynomials

We now describe a global approximation on the interval *w* ∈ [0, 1] that uses the Bernstein polynomials originally constructed by S. Bernstein [5] in 1912 and used to prove Weierstrass approximation theorem [6]. Given a function *f* (*x*), defined on the closed interval [0, 1], *the Bernstein polynomial of degree d of the function f* (*x*) is defined by:

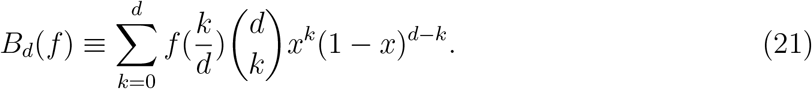

The expressions 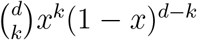 are called the Bernstein basis functions, the first seven of which (as well as many other interesting properties of them) are shown in figure 4 of [8]. It is proven that if *f* is continuous on [0, 1], then *B*_*d*_(*f*) converges uniformly to *f* on [0, 1] as *d* → ∞ with an error bound:

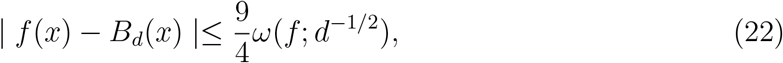

where the modulus of continuity [6] *ω*(*f, d*^−1*/*2^) is defined by

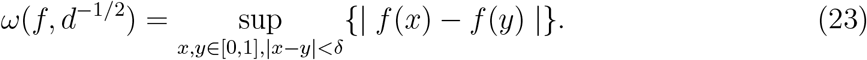

It is easy to see from formula (21) that *B*_*d*_(*f*) is a weighted linear combination of products of polynomials *x*^*k*^(1 − *x*)^*d*−*k*^, and it requires us to know *d* + 1 function values at discrete points 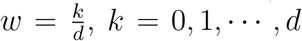, that equally partition the interval, [0, 1], into *d* subintervals. By contrast, the weak selection local approximation (15) or Taylor expansion approximations [3] are based on the stronger differentiability condition of 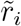 but at only one location *w* = 0. Tailoring the formula (21) for our use, we obtain the Bernstein approximation formula for the rate of evolution *r*_*i*_ as a function of the selection parameter *w* ∈ [0, 1]:

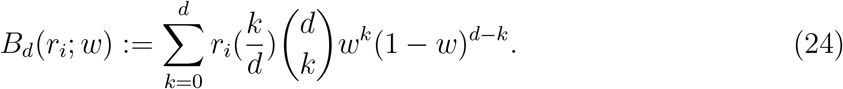

The Bernstein polynomial then becomes a global uniform approximation as long as *r*_*i*_ is continuous.

### Lemma 1.

*Given a two-player game for a fixed N, the Bernstein polynomial B*_*d*_ *of degree d uniformly converges to the rate of evolution r*_*i*_ *for all i* ∈ {1, 2, …, *N* − 1} *if the following inequalities are satisfied*

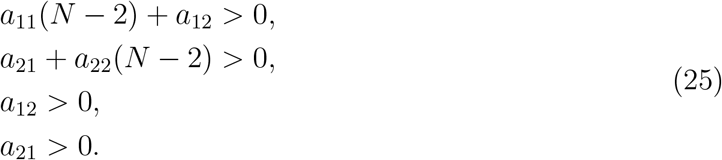

This sufficient condition is equivalent to 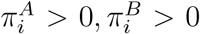 for all *i* = 1, 2, …, *N* − 1, which implies 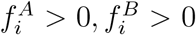 for all *w* ∈ [0, 1] taking into account that these are linear functions in *w*. Then, transition probabilities 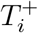 and 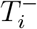 become positive, hence so is their ratio, eventually making *ρ*_*i*_ continuous in *w*.

It is clear that the Bernstein approximation *B*_*d*_ is a polynomial of degree less than or equal to *d* and passes through the two points (0, *f* (0)) = (0, 1) and (1, *f* (1)) for every *d*. Its definition requires *d* + 1 exact functions values 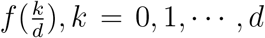. In particular, *B*_1_ is just the line segment joining the two end points of the curve *f* at 0 and 1. Moreover, the Bernstein approximation can be adjusted so as to approximate a function *f* defined on any closed interval [*a, b*] by scaling:

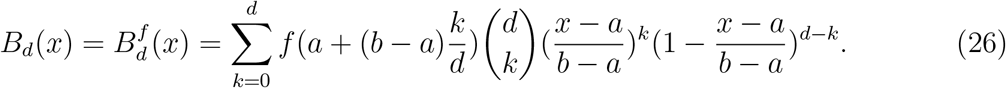

Thus whenever the interval of selection *w* of interest is narrowed, the Bernstein polynomial will be accordingly taken using the equation (26). When it is restricted onto the weak selection interval including the origin where the Taylor approximation gives a valid approximation, the global Bernstein approximation can be compared to this local (truncated) Taylor approximation. Differentiability at the origin is needed to construct this local pointwise Taylor approximation, while continuity is the main condition to form a uniform global Bernstein approximation. In fact, unlike the necessity of *d* + 1 original function values at different points in domain for the Bernstein approximation, *d* + 1 values of derivative functions *f* ^(*k*)^, *k* = 0, 1, …, *d* at the origin must be evaluated for a local Taylor polynomial of degree *d*.

To be clear, to actually use the formula, we need to pre-compute 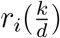 from the exact formula (14) at *d* + 1 grid points. The benefit of doing this is that we are then guaranteed that *B*_*d*_(*r*_*i*_; *w*) converges uniformly to *r*_*i*_ as *d* → ∞ throughout the entire region *w* ∈ [0, 1]. We show in figure 2 the Bernstein approximations to the rate of evolution *r*_1_ for *N* = 10, 100, 1000, 10, 000 throughout the full selection interval *w* ∈ [0, 1] as the degree increases. Note the formation of a boundary layer (non-uniformity) in the weak selection (left) regime. Also shown in the figures is the exact rate of evolution to which the Bernstein approximations converge. The exact rate of evolution curve is particularly interesting in that its shape depends very much on *N*, but as *d* increases, the finite-degree approximations converge to the exact (numerical) solution throughout the entire interval. The size of the errors between the Bernstein approximations and the exact formula is shown in figure 3. Notice that for any fixed value of *d*, the errors are largest in the weak selection (boundary layer) regime. Because of this, it is clear the Bernstein polynomial representation is particularly efficient in the strong selection regime *w* ∼ 1. To obtain high accuracy of the approximation in the other regimes would require more Bernstein modes. See [8] for more insight into convergence rates associated with using Bernstein polynomials to approximate functions.

**Figure 2:**
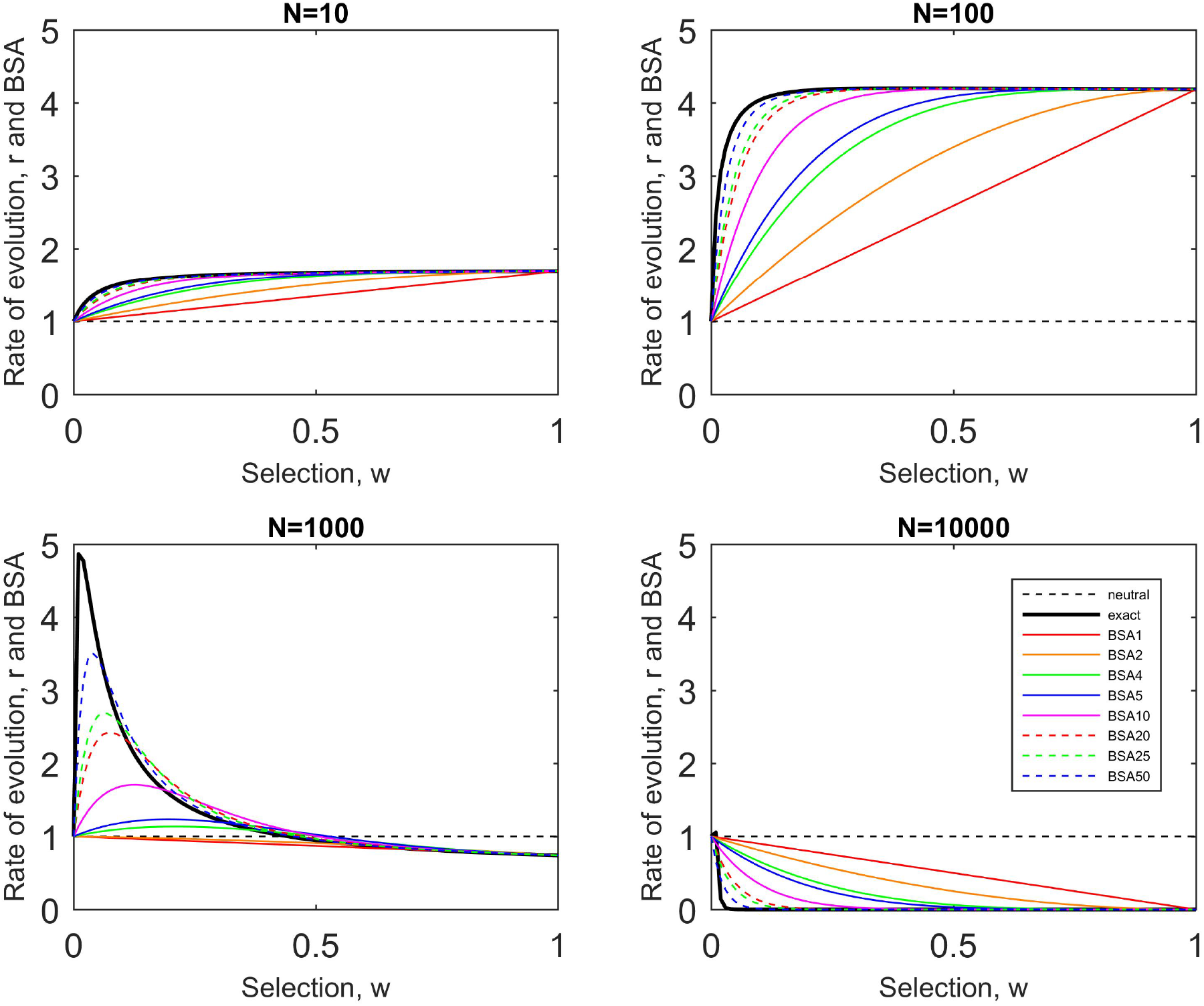
The Bernstein approximations over [0,1]. The Bernstein approximation *B*_*d*_(lines in colors) of various degrees of 1, 2, 4, 5, 10, 20, 25, 50 show uniform convergence to the rate of evolution *r*_1_ = *Nρ*_1_(black solid line) for each *N* = 10, 100, 1, 000 and 10, 000 as *d* increases. Note the boundary layer at the weak selection left edge. We use *a*_21_ = 5 *> a*_11_ = 3 *> a*_22_ = 1 *> a*_12_ = 0 for a repeated Prisoner’s Dilemma game with *n* = 10. The rate of evolution *r*_1_ for the neutral selection (black dashed line) is also given as a comparison.

**Figure 3:**
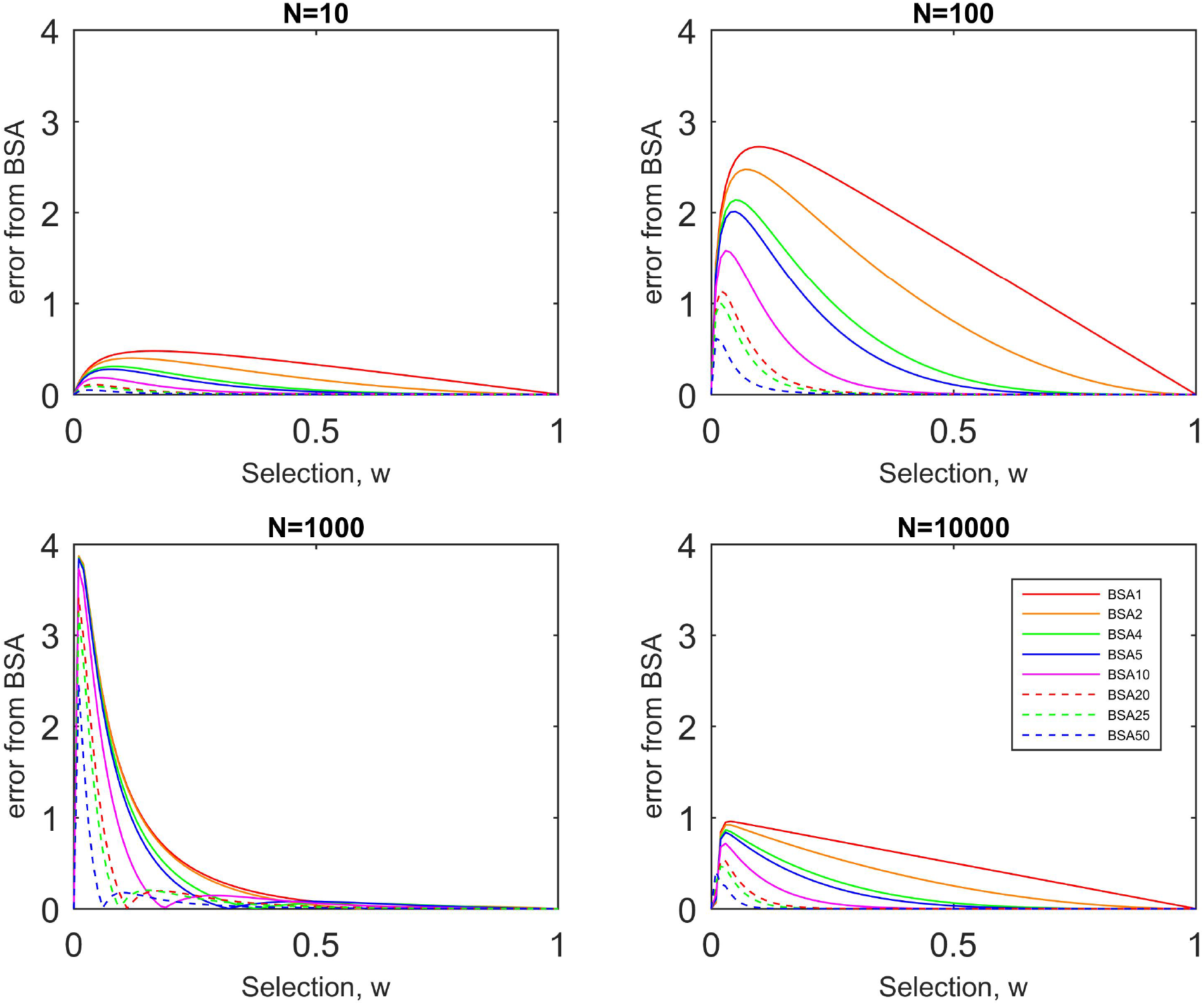
The Errors from the Bernstein approximations over [0,1]. We use *a*_21_ = 5 *> a*_11_ = 3 *> a*_22_ = 1 *> a*_12_ = 0 for a repeated Prisoner’s Dilemma game with *n* = 10. For any fixed *d*, errors are largest at the weak selection left edge and smallest at the strong selection right edge.

## 4. Discussion

The use of Bernstein polynomials as a basis set for approximating curves has a long and illustrious history, some of which is described in [8]. Because of the relatively slow convergence properties of the approximants, the method was not heavily developed as a practical tool until the advent of computers. Its current use (along with methods due to Bezier) is mostly in exploiting computers to interactively design parametric curves and surfaces because of the many insights the Bernstein coefficients provide on the properties of these sometimes complex objects. For our purposes, we use them as approximants to construct a uniformly valid approximation to a function defined via a stochastic process through and outside of a boundary layer region [4], which to our knowledge is novel. While we do not expect the approximation formula (24) to provide a practical substitute for the widely used formula (15) in the weak selection regime, we can imagine its use in problems where selection is stronger, or where it is computationally impractical to use the exact formula (14) over a range of selection parameter regimes. The approximation method we propose only requires the computation at *d*+1 evenly spaced grid points, and then we inherit (for free) all of the well known convergence properties of the Bernstein approximants [6, 8].

## 5. Conclusion

In summary, the Bernstein polynomial formula, *B*_*d*_(*r*_*i*_), (eqn (24)) plays the role of a global approximation to the fixation probability *ρ*_*i*_ and rate of evolution, *r*_*i*_ = *Nρ*_*i*_, on the full interval *w* ∈ [0, 1] for all *N* despite the presence of a boundary layer in the weak selection regime. Its performance improves as *d* increases for each *N* so that the error can be reduced as much as desired, and the major refinement is achieved for strong selection. However, the approximation needs larger values of *d* (degree) to gain the same accuracy in the weak selection range as the strong selection range, especially near *w* = 0, where it produces a relatively larger error because of the shape of the curve as a function of *N*. Despite the larger error near *w* = 0, this error is overall smaller for either a small or a large population size, as shown in Figure 3(a) and Figure 3(d). On the other hand, the error remains relatively large near *w* = 0 (even with a higher degree) for intermediate values of *N* as shown in Figure 3(c). In this weak selection range, there are several simple superior alternative formulas one might invoke as discussed earlier. It is probably in the intermediate and strong selection range that our Bernstein polynomial formulas might prove to be most useful.

## Acknowledgments

We gratefully acknowledge support from the Army Research Office MURI Award #W911NF1910269 (2019-2024) as well as useful conversations and suggestions from Prof. Kukavica and Prof. Ziane.

